# Characterizing higher order structures of chromatin in human cells

**DOI:** 10.1101/267856

**Authors:** Uwe Schwartz, Attila Németh, Sarah Diermeier, Josef Exler, Stefan Hansch, Rodrigo Maldonado, Leonhard Heizinger, Rainer Merkl, Gernot Läengst

## Abstract

Packaging of DNA into chromatin regulates DNA accessibility and, consequently, all DNA-dependent processes, such as transcription, recombination, repair, and replication. The nucleosome is the basic packaging unit of DNA forming arrays that are suggested, by biochemical studies, to fold hierarchically into ordered higher-order structures of chromatin. This defined organization of chromatin has been recently questioned using microscopy techniques, proposing a rather irregular structure. To gain more insight into the principles of chromatin organization, we applied an *in situ* differential MNase-seq strategy and analyzed *in silico* the results of complete and partial digestions of human chromatin. We investigated whether different levels of chromatin packaging exist in the cell. Thus, we assessed the accessibility of chromatin within distinct domains of kb to Mb genomic regions by utilizing statistical data analyses and computer modelling. We found no difference in the degree of compaction between domains of euchromatin and heterochromatin or between other sequence and epigenomic features of chromatin. Thus, our data suggests the absence of differentially compacted domains of higher-order structures of chromatin. Moreover, we identified only local structural changes, with individual hyper-accessible nucleosomes surrounding regulatory elements, such as enhancers and transcription start sites. The regulatory sites per se are occupied with structurally altered nucleosomes, exhibiting increased MNase sensitivity. Our findings provide biochemical evidence that supports an irregular model of large-scale chromatin organization.

## Introduction

The sequence-specific binding of proteins to DNA determines the activity of DNA-dependent processes, such as transcription, replication, repair and others, regulating cellular fate. However, nuclear DNA is packaged into chromatin, a nucleoprotein structure that restricts the access of specific DNA binding proteins. As a first level of compaction, DNA segments of 147 bp are wrapped in 1.7 left-handed turns around histone octamers forming the nucleosome core, each having a diameter of 11 nm. Thus, the human genome consists of approximately 30 million nucleosome cores, which are separated by DNA linkers whose length is cell-type specific and ranges between 20 to 75 bp (Prunell and Kornberg 1982; Compton et al. 1976; Woodcock and Ghosh 2010). These cores are the building blocks for higher levels of compaction and are assumed to fold at an intermediate level into fibres of 30, 120, 300, and 700 nm diameter, which ultimately constitute the mitotic chromosome (Woodcock and Ghosh 2010). This textbook model of hierarchical folding is based on the analysis of *in vitro* reconstituted chromatin and on chromatin extracted from permeabilized cells. The latest *in vitro* studies propose two alternative models for the 30 nm fibre: the one-start solenoid (Robinson et al. 2006) and the two-start zig-zag with approximately five to six nucleosomes per 11 nm of fibre (Schalch et al. 2005; Song et al. 2014).

However, existence of the 30 nm fibre and additional levels of chromatin compaction *in vivo* remains a controversial topic. Contrary to the textbook model, compact structures have been observed in terminally differentiated cells and specialized cells such as starfish sperm but not in proliferating cells (Horowitz et al. 1994; Scheffer et al. 2011; Fussner et al. 2011; Ricci et al. 2015). Additionally, a further series of experiments suggests that nucleosomes are highly interdigitated and do not form regular 30 nm fibres but irregular folded structures. These findings are best described by a polymer melt model (Joti et al. 2012; Chen et al. 2016a; Eltsov et al. 2008; Fussner et al. 2012; Hsieh et al. 2015; Nozaki et al. 2017; Ou et al. 2017). However, alternative higher-order structures, which are incompatible with the polymer melt model, have been described for metaphase and interphase chromosomes. Among them is the “rope flaking” model, having the nucleosomal arrays looped without self-crossing (Grigoryev et al. 2016). Such an organization would explain the release of several hundred kb-long chromatin loops after gentle lysis of the metaphase chromosomes (Marsden and Laemmli 1979). This model is supported by nuclease and topoisomerase II accessibility assays, which suggested that higher-order structures of chromatin are organized into 50 kb domains that form more compacted structures of 300 kb (Filipski et al. 1990). Taken together, these findings indicate that chromatin organization is still an enigma.

The degree of chromatin compaction is also correlated with gene transcription and is thought to impact the regulation of DNA-dependent processes. In paradigmatic studies, the sedimentation of the β–globin gene was monitored in sucrose gradients. Compared to bulk chromatin, a slower sedimentation was observed for the active gene, suggesting open chromatin (Caplan et al. 1987). Indeed, gene-rich domains are enriched in de-compacted chromatin regions that are maintained accessible by actively transcribing polymerases and the altered degree of DNA supercoiling within these domains (Gilbert et al. 2004; Naughton et al. 2013). Still, there is microscopic evidence that actively transcribed genes exist in a chromatin structure that is approximately 25 times more compact than the nucleosomal array, and only a 1.5- to 3-fold extension of the compacted fibre is observed upon transcriptional activation (Hu et al. 2009). Whether these changes in chromatin compaction are associated with a change of chromatin density or a loss of hierarchical packaging is not known.

High-throughput sequencing technologies allow the precise mapping of nucleosome positions throughout the genome and can reveal dynamic changes of nucleosome positions depending on the cell types or activating signals (Schones et al. 2008; Teif et al. 2012; Moshkin et al. 2012; Schones 2011; Diermeier et al. 2014). Recent studies have demonstrated disease-dependent differences in nucleosome positions in tissues and cell lines, suggesting an important role of chromatin architecture on cell fate (Valouev et al. 2011). However, these methods fail to elucidate the higher-order structures of chromatin. A recent study combined the Hi-C approach with chromatin fragmentation by micrococcal nuclease (MNase) in budding yeast (Hsieh et al. 2015). This approach allowed for chromatin analysis in the range of 200 bp – 4 kb, which would be suitable to address the organization of the 30 nm fibres. But, this study did not analyze global chromatin organization and suggested a variety of local folding motifs. Chromatin fragmentation using varying concentrations of MNase was used to reveal local changes in chromatin accessibility, and not focusing on the organization of higher order structures of chromatin (Chereji et al. 2015; Henikoff et al. 2011) (Mieczkowski et al. 2016).

We devised a new strategy to analyze differential chromatin accessibility from nucleosome resolution up to 1 Mb-size domains by comparing partial (low concentration) and complete (high concentration) MNase digestions of chromatin *in situ*. We generated high-coverage chromatin accessibility maps for HeLa cells and identified differential enrichment of high and low MNase-associated DNA fragments on large domains. Our systematic analysis revealed that coverage variations in the datasets show only sequence modulations, and therefore no distinct higher-order structures of chromatin are detectable in cells. Furthermore, we identified local hyper-accessible sites coinciding with active gene regulatory elements. Hyper-accessible sites are associated with MNase-sensitive nucleosomes, which are lost in high MNase but maintained in low MNase digestions. Thus, this analysis gives new insights into chromatin organization and challenges existing models of higher-order structures of chromatin.

## Results

### The rationale of the differential MNase-seq strategy

If chromatin forms distinct levels of higher-order structures, whose compaction increases from the open and active state to the repressed and inactive state, it follows that DNA accessibility must decrease with increasing compaction. As a consequence, we expect that DNA hydrolysis would first release accessible chromatin domains, followed by the release of compact chromatin domains at higher endonuclease concentrations. We used micrococcal nuclease (MNase), an endonuclease that preferentially hydrolyses DNA in the linker region between nucleosomes (Noll et al. 1975) as a tool to map chromatin accessibility. HeLa cells were permeabilized and incubated with increasing concentrations of MNase for 90 seconds. The DNA was purified and analyzed on agarose gels, showing increasing fragmentation of chromatin with higher MNase concentrations (Fig. 1A). Our lowest MNase concentration gave rise to a nucleosome ladder and even to mono-nucleosomal DNA. These findings are in full agreement with the hierarchical model of chromatin structure. Thus, we propose that at low-MNase conditions nucleosomes from open and accessible chromatin structures are released preferentially, whereas compact and inaccessible chromatin structures - which are more resistant -are only released at high-MNase conditions.

**Figure 1.**
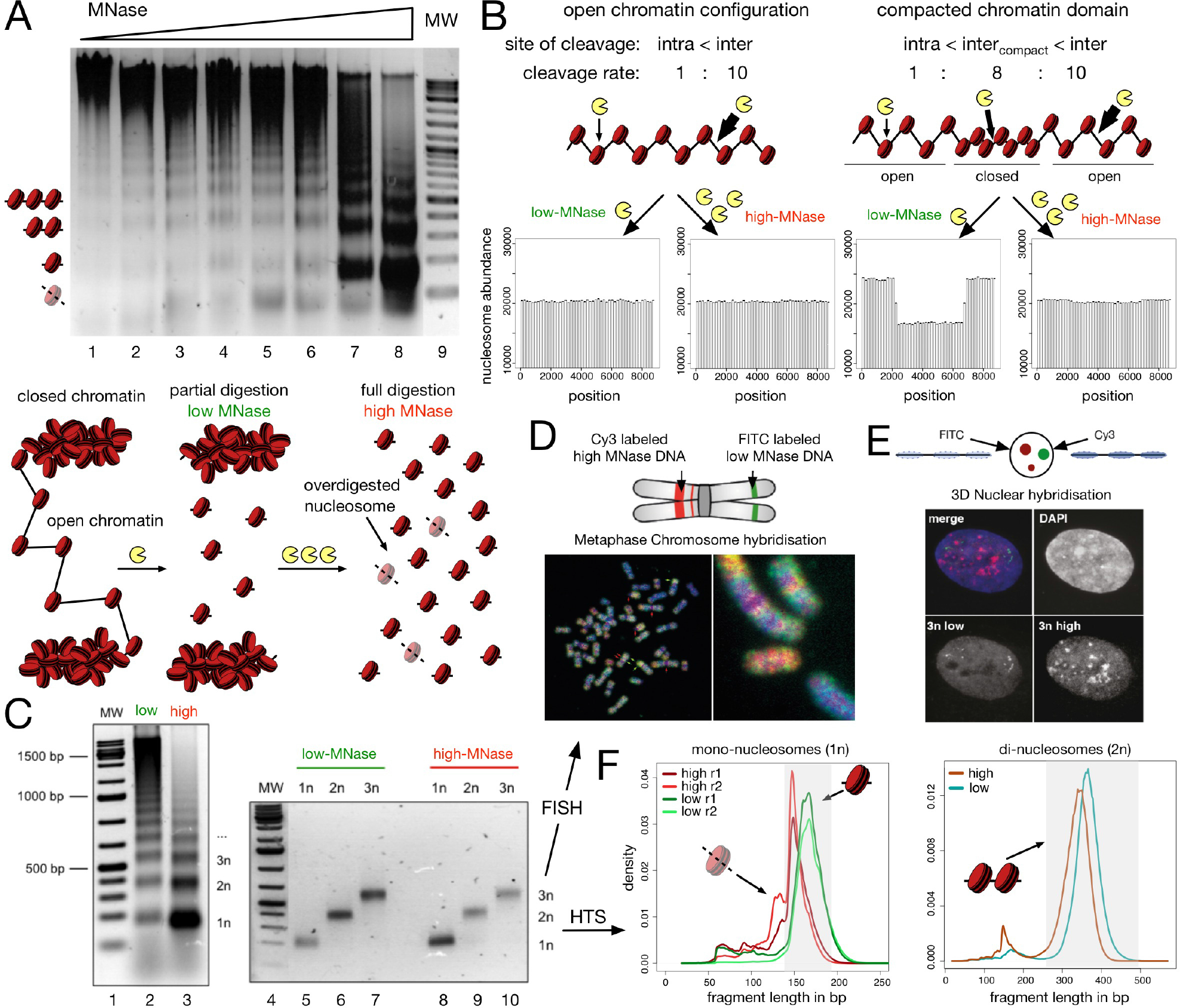
Isolation of nucleosomal DNA after differential MNase hydrolysis. (A) Nucleosomal DNA ladder of HeLa cells incubated with increasing concentrations of MNase. The agarose gel is showing the analysis of the purified DNA after MNase hydrolysis. The molecular weight marker is shown in lane 9 and the positions of subnucleosomal, mono-, di- and tri-nucleosomal DNA are indicated. The scheme below illustrates the hypothetical results of a partial and full hydrolysis of differentially accessible chromatin domains. (B) Simulation of low- and high-MNase digestions (see Methods for details) of an equally accessible linker DNA with *p*_cut_(*linker*) = constant (left panel) and of a nucleosomal array containing a central compacted chromatin domain with reduced linker accessibility *p*_cut_(*linker*, *compact*) = 0.8 · *p*_cut_(*linker*), (right panel). The relative sensitivity of intra-nucleosomal (intra) and inter-nucleosomal (inter) cleavage is indicated. (C) Large scale low- and high-MNase hydrolysis of chromatin (lanes 2 and 3) and subsequent isolation of nucleosomal DNA corresponding to the mono-, di-, and tri-nucleosomes (lanes 5 to 10). (D, E) 2D and 3D FISH experiments with isolated tri-nucleosomal DNA. Metaphase FISH (D) and 3D FISH (E) were performed with Cy3/FITC labelled tri-nucleosomal DNA derived from high-/low-MNase digestions, as indicated. (F) Low- ad high-digested mono- and di-nucleosomal DNA were subjected to library preparation and high throughput sequencing. The fragment size distributions of the mapped paired-end reads for mono- (left panel) and di-nucleosomal DNA (right panel) are shown. The shaded boxes highlight the fraction of isolated mono-nucleosomal (140 - 200 bp) and di-nucleosomal (250 - 500 bp) fragments.

### An *in silico* model for the differential MNase-seq strategy

To assess our initial assumption, we designed an *in silico* model that allowed us to simulate the effect of differential MNase hydrolysis on a nucleosomal array of 50 nucleosomes (Fig. 1B). We chose a stochastic approach and assigned different probabilities *p*_*cut*_(.) for MNase cleavage of DNA to different regions of the DNA (Fig. S1A). To begin, we defined two different regions and assigned *p*_*cut*_(*linker*) to DNA linker regions and *p*_*cut*_(*nuc*) to DNA covered by nucleosomes. The values *p*_*cut*_(*linker*) = 10 · *p*_*cut*_(*nuc*) were deduced from previous MNase footprinting studies (Cockell et al. 1983). Altogether, we modelled 101 cleavage sites, 50 intra-nucleosomal and 51 inter-nucleosomal DNA linker sites. According to the *p*_*cut*_(.) values, regions were chosen randomly, and if a region was selected by our algorithm, a cut was introduced by labelling the central nucleotide. At the end of the simulation, these labels were used to deduce the number and length of the fragments.

The characteristic difference of our partial (low-MNase concentration) and complete (high-MNase concentration) chromatin digestion conditions is reflected by the number of cutting events per template. In accordance with our experimental conditions, we assigned 20 double stranded cutting events for the low and 70 for the high MNase condition in all the computer simulations (Fig. S1B). The *in silico* MNase digestion of the nucleosomal array was iterated until 1 million mono-nucleosome-sized fragments were collected for each condition. Finally, the location and number of mono-nucleosomal fragments were determined and plotted. As expected, computer simulations modelling at least 20 cutting events resulted in histograms that resembled the nucleosome ladders of our experimental setup (Fig. S1B, S1C); these findings confirmed our model and choice of parameters.

We wanted to know how the open and compact chromatin domains would be represented in the high-and low-MNase digestion conditions. Modelling a chromatin domain with identical DNA linker accessibility (*p*_*cut*_(*linker*) = const) throughout the whole nucleosomal array resulted in indistinguishable accessibility patterns at varying MNase digestion conditions (Fig. 1B, left panel and Fig. S2A). However, if we modelled a compacted chromatin domain located at the centre of a chromatin fibre by means of 25 nucleosomes with reduced DNA linker accessibility (*p*_*cut*_(*linker*, *compact*) = 0.8 · *p*_*cut*_(*linker*), we observed striking differences between low and high digestion conditions (Fig. 1B, right panel). For high-MNase conditions, which are the standard experimental conditions to study nucleosome occupancy and cellular chromatin structure, modelling resulted in identical patterns. In contrast, strikingly reduced levels of nucleosomal DNA were predicted for the compacted chromatin domains, when we simulated low-MNase conditions (Fig. 1B, right panel). Note that this finding does not depend on the chosen parameters: The simulation congruently predicted differences between MNase conditions in the compacted chromatin domain, simulating 10 to 60% of array cleavage (Fig. S2B). Furthermore, the low-MNase condition sensitively detected alterations of the linker accessibility, whereas the high-MNase signal was unaffected (Fig. S2C). Taken together, the outcome of our simulation suggests that low-MNase conditions, in comparison to high-MNase conditions, should reveal differences in hierarchical chromatin organization.

### High- and low-MNase treated chromatin localizes to distinct nuclear compartments

To test our computer modelling experimentally and to annotate chromatin domains, we isolated the mono-, di- and tri-nucleosomal DNA fractions of low- and high-MNase treated cells for FISH (fluorescence *in situ* hybridization) and high-throughput sequencing analysis (Fig. 1C - F). First, we tested whether low- and high-MNase treatment released different fractions of the genome by analysing the tri-nucleosomal DNA in 2D and 3D FISH experiments. FISH is an ideal method to visualize differential genomic locations of DNA in the mega base pair 11scale by hybridization to metaphase chromosomes (2D FISH), as well as to monitor the compartmentalization of the DNA probes in the nucleus by retaining the 3D structure of the chromosomes in the cell nucleus (3D FISH). The tri-nucleosomal DNA isolated from high- and low-MNase conditions was fluorescently labelled with either Cy3 (high-MNase) or FITC (low-MNase) and was subsequently used as probes in 2D and 3D FISH (Fig. 1D and E). In 2D, the signals of the high- and low-MNase DNA probes partially overlapped, but large regions with distinct staining patterns could also be detected (Fig. 1D). As expected, we observed regions devoid of low-MNase-treated nucleosomal DNA within the centromeric regions of the chromosomes. Additionally, the 3D FISH experiment, maintaining the nuclear architecture in 3D, clearly showed a spatial separation of the chromatin regions extracted with either low- or high-MNase concentrations (Fig. 1E). We also performed Southern blot experiments, hybridizing the MNase-treated chromatin with the isolated tri-nucleosomal DNA from high- and low-MNase treatment (Fig. S3). Hybridizing the high- and low-MNase-treated tri-nucleosomal DNA to nucleosomal ladders showed indistinguishable hybridization patterns. This suggests that differential MNase treatment does not result in a global loss or enrichment of genomic fractions. Therefore, our results indicate that high- and low-MNase extractions do preferentially release different genomic regions, potentially exhibiting distinct DNA packaging properties and localizing to different nuclear compartments in the cell.

### MNase sequence preferences dominate differential chromatin fragmentation

To resolve and map the different chromatin domains at high resolution, we performed high-throughput sequencing with the isolated mono- and di-nucleosomal DNA, extracted with either high- or low-MNase concentrations. Plotting the length distribution of the paired-end sequencing reads showed the expected average DNA fragment length of ~150 bp for mono-nucleosomal DNA and ~340 bp for di-nucleosomal DNA (Fig. 1F). Nucleosomal DNA fragments released by low MNase digestions tend to be slightly larger due to incomplete hydrolysis of the linker DNA. Interestingly, the high-MNase-treated chromatin exhibited an additional fraction of sub-nucleosomal sized fragments, which may represent partially digested nucleosomes (Ishii et al. 2015). For our subsequent analyses, we extracted DNA fragments in the size range of 140 to 200 bp for the mono-nucleosomal DNA and 250 to 500 bp for the di-nucleosomal DNA, yielding in total 393 million annotated mono-nucleosomes and 75 million di-nucleosomes. The read depth allowed us to perform a thorough statistical analysis of genome coverage and subsequent chromatin domain analysis.

Mapping the sequence reads to the human genome (version hg19) was performed and an overview of mono-nucleosomal read distribution is exemplarily given for chromosome 6 (Fig. 2A) and the whole genome (Fig. S4A). The distribution of low- and high-MNase-treated di-nucleosomal DNA reads is shown in Figure S4B and C. In both cases, the high- and low-MNase-treated samples revealed a high coverage and a rather similar distribution across the genome. To monitor differential fragment enrichments with respect to low- and high-MNase treatment, the log fold-change in coverage was calculated in 1 Mb non-overlapping windows (Fig. 2A, Fig. S4). Interestingly, as in the FISH experiments, we observed clusters of large, Mb-sized domains, exhibiting either enrichment of high- or low-MNase-treated chromatin fractions. These findings suggest that chromatin fragmentation and the subsequent isolation of mono- and di-nucleosomes gave rise to an enrichment pattern that is specific to the applied MNase concentration.

**Figure 2.**
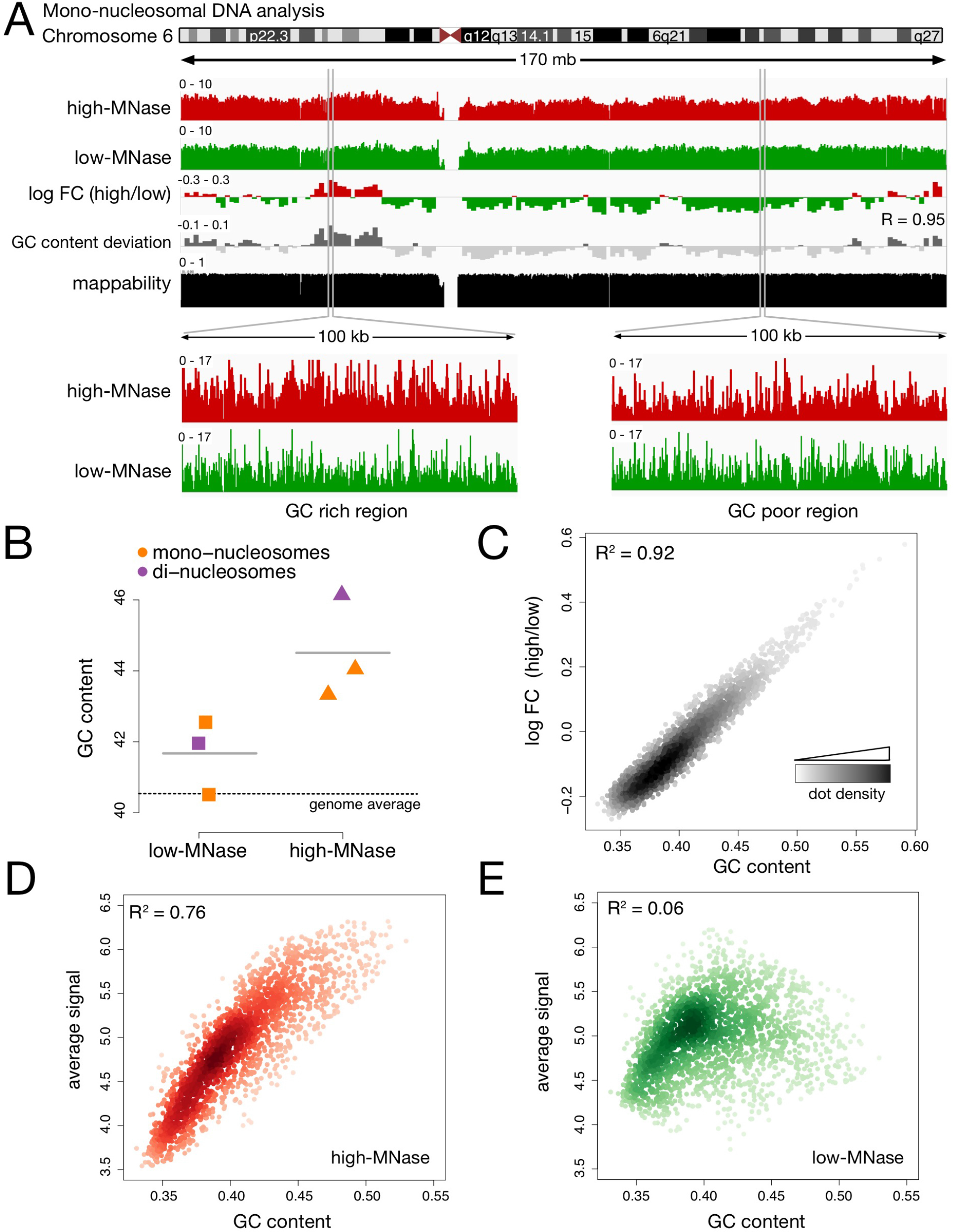
MNase sequence preferences dominate differential chromatin fragmentation. (A) Genome browser plot showing the nucleosome occupancy distribution along chromosome 6. Differences of high- and low-MNase digestions were calculated as log2 fold changes (log FC) of the average profile intensity in 1 Mb non-overlapping windows. GC content variation is displayed as the deviation from the genome wide average (40.5%). R is the Pearson correlation coefficient determined for the log FC (high/low) and the GC content deviation. The mappability track at the bottom illustrates how uniquely 100mer sequences align to a region of the genome (Derrien et al. 2012). (B) Average GC content of isolated mono- and di-nucleosomal fragments. The genome wide GC average is indicated by the dashed line. (C - E) Genome wide correlation of GC content und nucleosome occupancy profiles. Dots represent 1 Mb non-overlapping windows; the coefficient of determination (R^2^) refers to the result of a simple linear regression between GC content and the average signal of (C) log FC of high-MNase versus low-MNase, (D) high-MNase, (E) low-MNase.

MNase has a known sequence preference, preferentially hydrolyzing DNA at sites exhibiting A/T nucleotides (Hörz and Altenburger 1981; Dingwall et al. 1981; Flick et al. 1986; McGhee and Felsenfeld 1983; Ishii et al. 2015). Therefore, we analyzed the GC content of the isolated DNA fragments. Remarkably, genomic regions enriched in the high-MNase digestions showed an increased GC content, whereas the average GC content of the low-MNase-isolated fragments reflected the genome-wide average as indicated by the *p*-value = 0.03 of a paired t-test (Fig. 2B). To exclude the possibility that this is a specific characteristic of our experimental approach, we applied the same analysis to previously published data sets obtained from different organisms (Chereji et al. 2015; Ishii et al. 2015). Even though different chromatin extraction protocols have been used, including formaldehyde crosslinking and nuclei isolation steps, the same trend can be observed: The GC content of chromatin released with high-MNase concentrations is higher, whereas the GC content of low-MNase samples is generally close to the genome-wide GC distribution (Fig. S5). Furthermore, from those data sets we deduced additional controls following MNase digestion, such as Histone-ChIPs, and confirmed that this effect is not caused by a contamination of non-nucleosomal fragments in the low-MNase condition (Fig. S5).

To test whether the sequence preference of MNase would mask the identification of chromatin domains, we correlated the log fold-change of high- and low-MNase signals with the underlying GC content. Indeed, we observed a remarkably high correlation between GC content and differences in high- and low-MNase digestion (R^2^ = 0.92, Fig. 2C), indicating that the DNA sequence influences the differential chromatin fragmentation (Fig. 2A - C, Fig. S6A and B). The comparison of a GC-poor and a GC-rich genomic region revealed great variations of the high-MNase signals, yielding more reads in high-GC regions and less reads with a more variable read distribution in low-GC regions (Fig. 2A, lower panel). The genome-wide correlation analysis indicated a strong correlation between the signal under high-MNase conditions and the GC content (R^2^ = 0.76) explaining the varying signal intensities (Fig. 2D). In contrast, the low-MNase reads did not correlate with GC content (R^2^ = 0.06, Fig. 2E). This finding showed that the low-MNase digestion pattern is unaffected by the GC content, giving rise to a homogenous read distribution across the genome.

### Nucleosomes of GC-poor genomic regions are under-represented in high-, but not in low-MNase conditions

MNase-dependent sequence bias and the positioning and occupancy of nucleosomes have a strong effect on the dinucleotide repeats of the isolated DNA fragments. Therefore, we wanted to know whether these were equally well represented under low- and high-MNase conditions. For both low-and high-MNase conditions we could confirm two typical signatures: First, the overall dinucleotide frequencies within the nucleosomal DNA fragments showed the expected 10 bp phasing of G/C or A/T dinucleotides (Kaplan et al. 2009), confirming the proper isolation of nucleosomal DNA (Fig. 3A, Fig. S7). Additionally, we detected the preferential hydrolysis of DNA at A/T sites in isolated DNA fragments (Ishii et al. 2015) (Fig. S7B).

**Figure 3.**
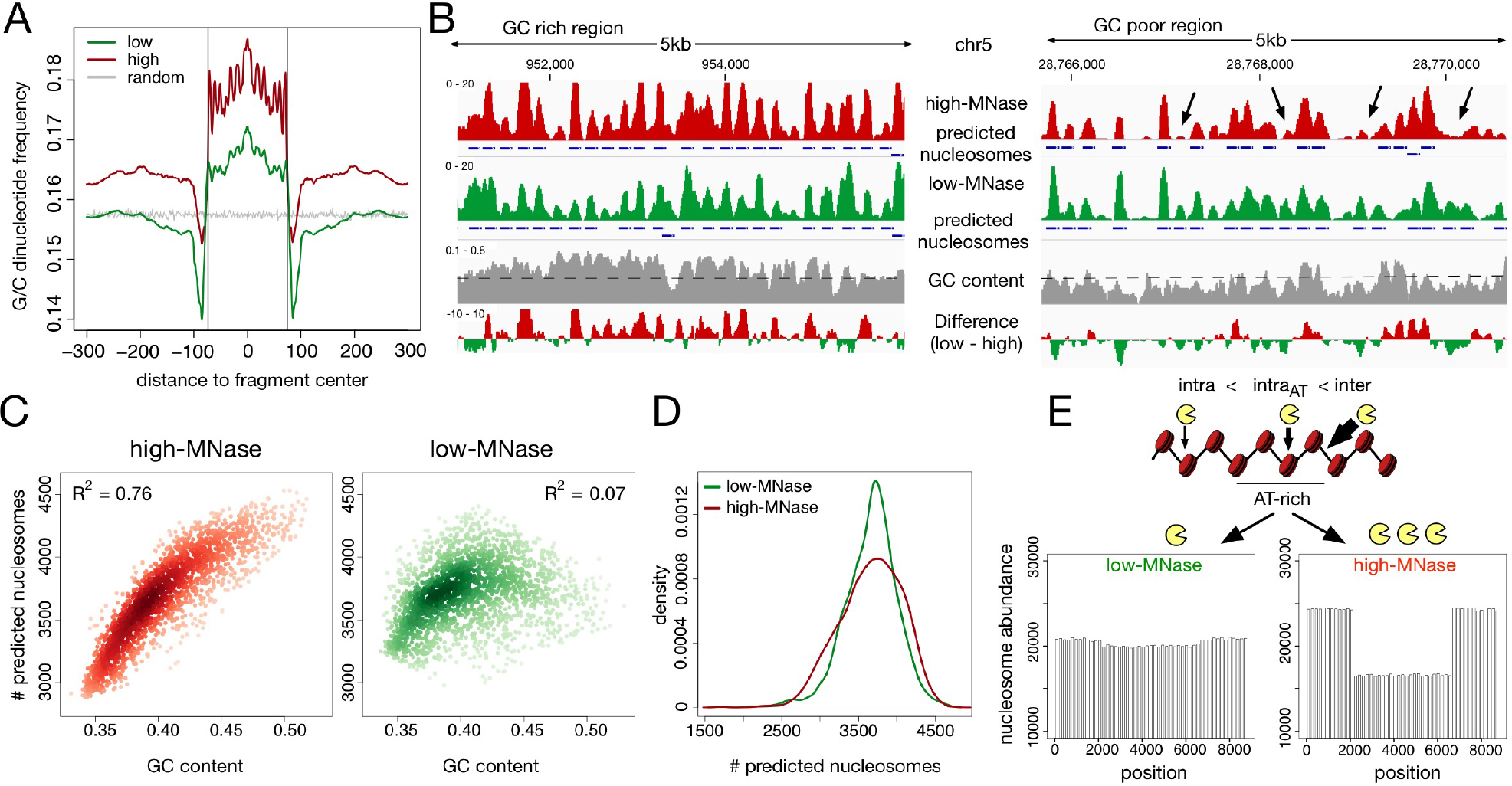
Nucleosomes in AT-rich regions are partially depleted in high-, but not in low-MNase conditions. (A) Average G/C-di-nucleotide frequency, deduced from GG, GC, CG, CC occurrences, centred on mapped mono-nucleosomal fragment midpoints. A random distribution (grey) was simulated on 1 million randomly generated fragments of 147 bp lengths. Vertical lines indicate nucleosome boundaries (± 73 bp). (B) Genome browser snapshot showing the GC bias of nucleosome annotation after high MNase digestions. Nucleosome positions, which have been called with the `DANPOS2` toolkit (Chen et al. 2015), are represented by blue boxes. Arrows indicate over-digested nucleosomes in high MNase condition. GC content was calculated in 50 bp windows with 10 bp sliding steps. The genome wide GC average is given by the dashed line. (C) Genome wide correlation of GC content and quantity of nucleosome prediction. Dots represent 1 Mb non-overlapping windows; (D) Kernel density plot of the nucleosome count in 1 Mb sized windows. (E) Simulation of low- and high-MNase digestions of a nucleosome strand exhibiting nucleosomes with variable MNase sensitivity. A higher intra-nucleosome cleavage probability *p*_*cut*_(*nuc*, *AT*+) was assigned to a stretch simulating an AT-rich region in the middle of the chromatin strand. Used cleavage probabilities: *p*_*cut*_(*nuc*, *AT*+) = 2 · *p*_*cut*_(*nuc*) and *p*_cut_(*linker*) = 10 · *p*_*cut*_(*nuc*).

However, in addition to the local A/T preference of MNase, we observed changes in nucleotide content within the nucleosome core sequence and neighboring regions that differ with the experimental condition. The nucleosomal DNA possessed decreased A/T dinucleotide levels and increased G/C content relative to the genomic levels in low-MNase conditions, and this trend became more prominent in high-MNase conditions (Fig. 3A, Fig. S7A). When using low-MNase conditions, the average G/C content in the linker DNA and surrounding regions matched the genome-wide dinucleotide distribution. In contrast, when using high-MNase conditions we observed an overall increase in the G/C dinucleotide content of the nucleosomal DNA and the surrounding genomic regions. This suggests that high-MNase treatment introduces the preferential selection of GC-rich fragments. This effect can be explained by an increased MNase sensitivity of nucleosomal core DNA reconstituted on AT-rich DNA, thereby depleting these nucleosomes from the pool (Fig. 2A, lower panel). Next, we addressed whether the observed GC bias in high-MNase digestions would influence the analysis of genome-wide nucleosome positions and occupancy. Therefore, we used the `DANPOS` package (Chen et al. 2013) to annotate nucleosome positions from our MNase-seq data. `DANPOS` identified and annotated fewer nucleosomes in GC-poor regions of high-MNase digestions than with low-MNase treatment (Fig. 3B). Among the low-MNase data, the overall number of predicted nucleosomes depended on the underlying GC content and nucleosome prediction was more homogenous throughout the genome (Fig. 3C and D). In line with the observed GC bias in the read distribution of the high-MNase data, we observed that nucleosome annotation is worse in the GC-poor regions.

So far, our computer simulation did not predict a variation in the high-MNase data (Fig. 1B), which indicated that a further refinement of our *in silico* approach was required. To model the preferential digestion of nucleosomes occupying AT-rich regions, we introduced the probability *p*_*cut*_(*nuc*, *AT*+) as *p*_*cut*_(*nuc*, *AT*+) = 2 · *p*_*cut*_(*nuc*). The outcome of this refined model resembled our *in vivo* observations at high-MNase-conditions (Fig. 3E). Now, domains with AT-rich DNA sequences were represented by a loss of extracted nucleosomes at high-MNase conditions. Interestingly, the refinement of the model did not alter nucleosome extractability in the low-MNase conditions. Note that this prediction does not depend on the chosen *p*_*cut*_(*nuc*, *AT*+) values and is robust even at higher nucleosome cleavage probabilities (Fig. S8). Our *in silico* results are in agreement with the experimental findings from low-MNase conditions which provide a more detailed representation of genome-wide nucleosome positions than the commonly used high-MNase concentrations.

The inherent GC bias could interfere with the analysis of domain architecture in the high-MNase conditions. In contrast, the above findings make plausible that low-MNase conditions allow for the identification of inaccessible domains from higher-order structures. To confirm this assumption, we compared the low-MNase data with the read distribution of sonicated chromatin, free of sequence bias. As shown in Figure S9A and B, the datasets are highly correlated (R = 0.8); this finding suggests that there are no genomic domains of differential DNA compaction on the mega base pair scale (Mb).

### Large domains of differential chromatin compaction are not detectable

Next, we addressed whether differential compaction states of chromatin would affect the MNase dependent extraction of DNA in genomic regions that are smaller than the Mb scale. Domains located in open or compacted chromatin configurations should modulate DNA accessibility, independent of the underlying GC content. Therefore, we used a sliding window approach and determined the correlation of the low- and high-MNase datasets with the GC content of the corresponding genome sequence. This analysis is based on the assumption that a fraction of the genome should escape the GC content correlation of the low-/high-MNase extraction, if differential chromatin packaging domains exist.

First, correlation was determined for a window size of 1 Mb, which corresponds to a domain of approximately 5000 nucleosomes. The very high values (R^2^ = 0.92) argue against differential chromatin domains of this size (Fig. 4A). A pronounced correlation (R^2^ > 0.44) persisted, even after gradually reducing the scanning window size down to 5 kb, which represents a domain of only 3 solenoidal turns and a total of approximately 25 nucleosomes. Even for a window size of 1 kb which is too small to represent a structural domain, we observed a moderate correlation (Fig. 4A, Fig. S10A).

**Figure 4.**
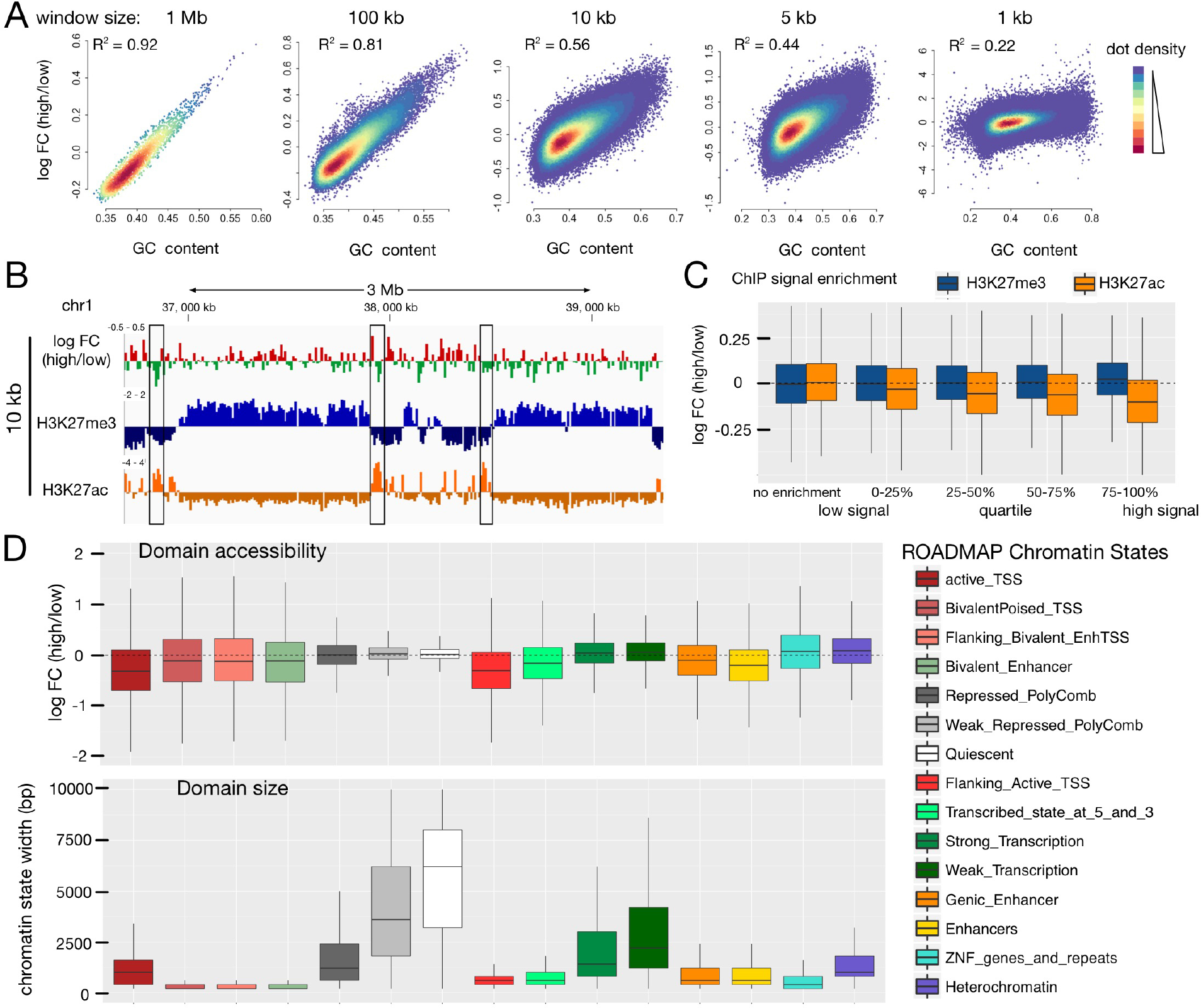
Comparison of chromatin accessibility within annotated chromatin regions. (A) Genome wide correlation of GC content and differential MNase signal. Dots represent non-overlapping windows of varying genome sizes (size given at the top). (B) Genome browser snapshot showing H3K27me3 (ENCODE, ENCFF958BAN) marked domains and local H3K27ac sites (ENCODE, ENCFF311EWS) highlighted by black rectangle compared to differential MNase digestions. log_2_ FC was calculated in 10 kb non-overlapping windows. (C) Correlation of H3K27me3 (blue) / H3K27ac (orange) enrichment over input with differential MNase digestion enrichment. The average signal was calculated in 10 kb non-overlapping windows spanning the whole genome. Windows were grouped on the basis of the ChIP enrichment signal into windows showing no ChIP enrichment (ChIP-signal < input signal) and windows enriched for the histone modification were further subdivided into quartiles of signal enrichment. (D) Differential MNase signal (upper panel) in annotated chromatin states (ROADMAP) compared to the size of the chromatin states (lower panel). (B-D) The MNase signal was normalized for the underlying GC-bias (see Fig. S11 and Methods for details).

To directly assess the compaction states of euchromatin and heterochromatin fractions, we then focussed on the annotated regions that can be identified by their post-translational histone modifications. However, due to the dominating GC bias in our analysis, we were not able to correlate differences of low-/high-MNase with histone modifications (Fig. S10). Therefore, we globally normalized the number of sequencing reads to the GC content of the underlying genomic region. Generally, euchromatin has a higher GC content than heterochromatic regions (Comings et al. 1975). Thus, we applied a locally weighted scatterplot normalization (LOESS, window size 250 bp) to correct for the genome-wide GC variation (Fig. S11). After normalization, the ratio of low- and high-MNase sequencing reads were no longer correlated with GC-content (Fig. S11, lower panel; R = 0.02), allowing us to screen for other features that might modulate DNA accessibility.

We first had a closer look at a heterochromatin locus on chromosome 1 that is heavily marked by histone H3K27me3. Visual inspection showed no tendency of particular high-/low-MNase ratio changes within this domain (Fig. 4B). This locus was not a special case and we observed no correlation of high-/low-MNase ratios with increasing histone H3K27me3 density for a genome-wide of heterochromatin (Fig. 4C, blue boxplots). In contrast, euchromatic marks such as H3K27ac are not spread over large genomic regions but are locally concentrated (Fig. 4B). The selected genomic region was strongly enriched for H3K27ac modifications, overlapping with a domain preferentially enriched in the low MNase dataset, indicating locally increased accessibility of MNase (Fig. 4B, boxes). This is even true on the genomic scale: With higher densities of the H3K27ac mark, we observed an increased accessibility towards MNase hydrolysis (Fig. 4C). This suggests that only highly modified sites like active enhancers show an alteration in the chromatin structure.

To elucidate features that might affect the accessibility of chromatin, we statistically analyzed the annotated chromatin states and their genomic regions from the ROADMAP project (Roadmap Epigenomics Consortium et al. 2015) (Fig. 4D). Therefore, the high-/low-MNase ratio in functionally defined domains of the genome was calculated. Box-plots with negative log FC values are indicative of elements located in more accessible chromatin. These were active Transcription Start Sites (TSS), bivalent poised TSSs, enhancers and regions flanking the active TSSs and transcribed regions. Analysis of domain sizes revealed that these regions ranged from 200 to 1500 bp and showed that chromatin accessibility is regulated at a local scale. In contrast, no compacted genomic regions with decreased MNase sensitivity were detected. The heterochromatic and transcriptionally repressed regions of the genome, encompassing domains of 1000 to 10000 bp, showed similar MNase accessibility at both low and high enzyme concentrations. Taken together, these analyses, which were deduced from several, large (> 1.5 kb) genomic regions, suggest a homogeneous accessibility of chromatin that does not depend on histone modification patterns or functional domains of the genome. As a consequence, we postulate that domains of differential DNA packaging do not exist.

### Chromatin accessibility is modulated on local scale

As we observed differential chromatin packaging at the local scale, we performed a detailed analysis of the low-/high-MNase data sets at the nucleosomal level. Sliding windows as small as 1 kb exhibited a decreased GC correlation (Fig. 4A, Fig. S10B and C). Remarkably, sites showing an enrichment in the low-MNase fraction coincided with DNase I HS sites and/or active histone marks as H3K27ac near actively transcribed genes. Visual inspection suggested an enrichment of nucleosomes released under low-MNase conditions at those sites (Fig. 5A, Fig. S10B and C), suggesting the existence of local hyper-accessible sites.

**Figure 5.**
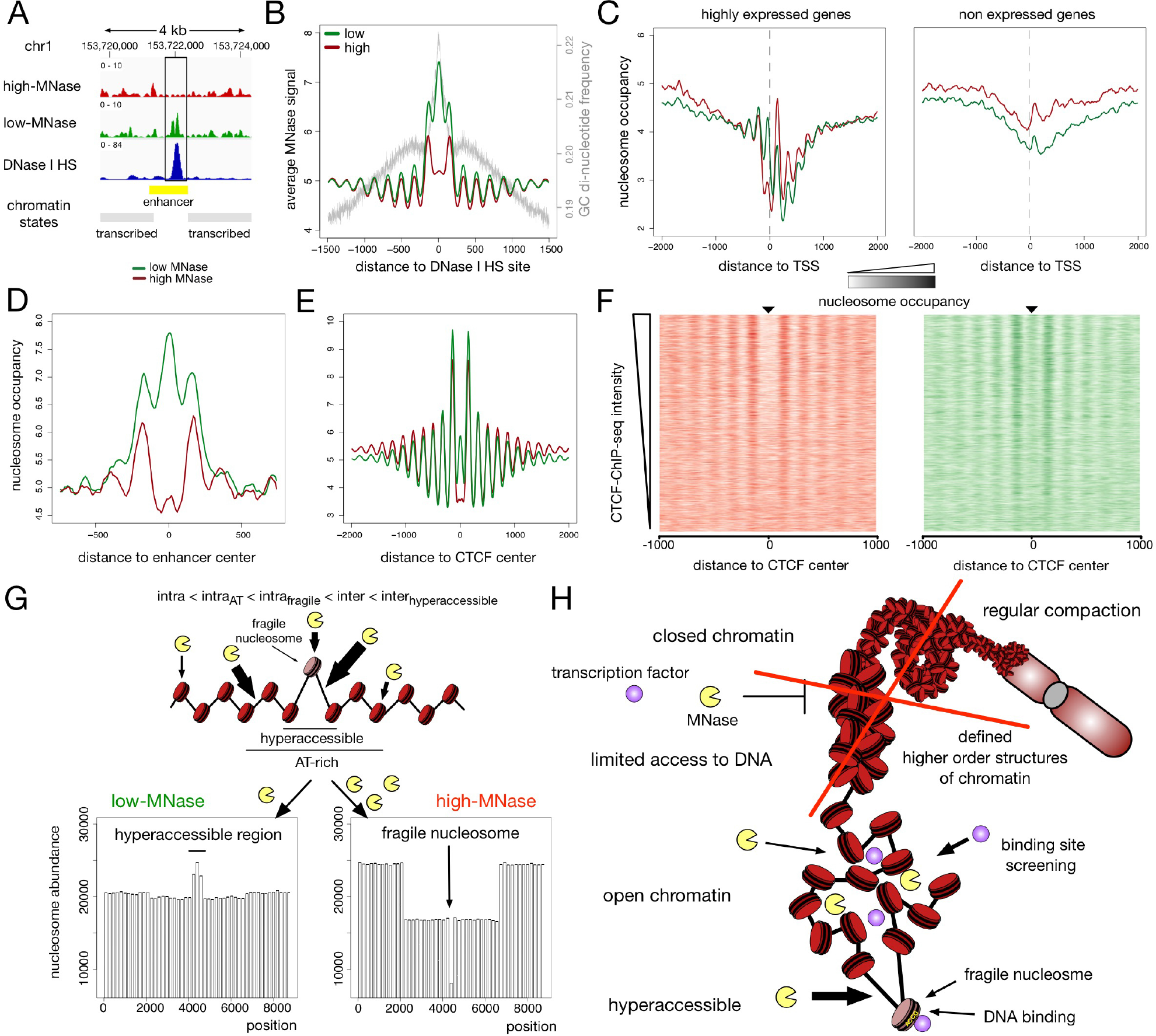
Differences of low- and high-MNase analysis at local sites. (A) Genome browser snapshot around an accessible enhancer marked by DNase I HS peak (ENCODE, ENCFF567PRQ). (B - E) Averaged MNase-seq profiles around: (B) DNase I HS sites (ENCODE, ENCFF692NCU); average G/C - di-nucleotide content is indicated by the grey line. (C) TSS of highly expressed genes (left panel) and not expressed genes (right panel). FPKM values determined by RNA-Seq analysis (ENCODE, ENCFF000DNW) were used to estimate the level of gene expression. Genes not detected in RNA-Seq analysis (FPKM = 0) were considered as not expressed and those genes showing the 25% highest FPKM values were considered as highly expressed. (D) Active enhancer sites in HeLa cells as defined in (Chen et al. 2016b). (E) CTCF sites (ENCODE, ENCFF002DCW). (F) Heatmaps of high-MNase (left panel) and low-MNase (right panel) are centred on CTCF binding sites and sorted in descending order of ChIP-Seq density at detected CTCF sites. (G) Simulation of low- and high-MNase digestions of a nucleosome array exhibiting nucleosomes with variable MNase sensitivity and hyper-accessible DNA sites. A higher intra-nucleosome cleavage probability *p*_*cut*_(*nuc*, *AT*+) was assigned to a domain, simulating an AT-rich region in the middle of the chromatin strand. In the centre of the nucleosomal array an hyper-accessible site was introduced, representing a fragile nucleosome with markedly increased intra-nucleosome cleavage probability *p*_*cut*_(*nuc*, *fragile*), and the two adjacent DNA linker sites were assigned with enhanced inter-nucleosomal cleavage probability *p*_*cut*_(*linker*, *hyperaccessible*). Chosen cleavage probabilities: *p*_*cut*_(*nuc*, *AT*+) = 2 · *p*_*cut*_(*nuc*), *p*_*cut*_(*nuc, fragile*) = 4 · *p*_*cut*_(*nuc*), *p_cut_(linker)* = 10 · *p*_*cut*_(*nuc*), *p_cut_(linker, hyper-accessible)* = 12 · *p*_*cut*_(*nuc*).

Next, we correlated the nucleosome profile of either high- or low-MNase digestions with annotated DNase I HS in HeLa cells from the ENCODE project (Roadmap Epigenomics Consortium et al. 2015). Though the GC content at DNase I HS is higher than the genome-wide average, our analysis showed an enrichment of the low-MNase fractions, indicating that nucleosomes at these sites adopt an open chromatin configuration (Fig. 5A and B). Interestingly, only nucleosomes coinciding with or adjacent to the DNase I HS showed a significant enrichment, suggesting local modulation of chromatin accessibility. In addition, the MNase sequence bias within the high-MNase dataset would result in an underestimation of accessible nucleosome enrichment. Therefore, we corrected GC enrichment by using a LOESS normalization as described above (see above and Methods for details, Fig. S11). We applied the `Dpeak` script from the `DANPOS2` toolkit to the normalized data in order to identify differentially enriched sites in the low- and high-MNase fractions. In the genome wide analysis, 36433 sites revealed a significant enrichment in low-MNase nucleosomes and were termed as MNase hyper-accessible sites. In addition, we identified 35454 sites with significant enrichment in high-MNase nucleosomes, termed MNase inaccessible sites (Fig. S12A and B). In contrast to MNase inaccessible sites, MNase hyper-accessible sites coincide with the annotated DNase I HS (*p*-value < 2.2·10^16^ Fisher’s exact test, Fig. S12B). Although we observed only a partial overlap, DNase I HS coinciding with MNase hyper-accessible sites exhibited a high signal intensity in DNase I HS assays (Fig. S11C). Furthermore, MNase hyper-accessible sites were enriched at gene regulatory elements, such as active TSS and predominantly enhancers, indicating a link between local chromatin accessibility and gene regulation (Fig. S12D). In summary, our data suggests that the nucleosomal array structure is exposed only locally, potentially allowing for the binding of regulatory factors.

### MNase-sensitive nucleosomes occupy hyper-accessible sites

A recent study using a chemical approach to map nucleosomes in mouse embryonic stem cells identified histone binding at regulatory sites, which have been suggested to be nucleosome-free in genome-wide MNase-seq studies (Voong et al. 2016). In agreement with this chemical mapping approach, we identified well-positioned nucleosomes upstream of the active TSS using the low-MNase conditions (Fig. 5C). Presumably, these nucleosomes have escaped previous MNase-seq studies which used MNase conditions that we describe as high-MNase.

Accordingly, the centres of active enhancers in HeLa cells were depleted of nucleosomes in the high-MNase fraction, whereas in low-MNase fractions these nucleosomes were markedly enriched (Fig. 5D). Even directly at CTCF-bound sites we detected MNase-sensitive nucleosomes, which were preferentially degraded in high-MNase conditions (Fig. 5E). In contrast to a model, where CTCF and nucleosomes compete for DNA binding, we observed the highest nucleosome enrichment in low-MNase at sites exhibiting strong CTCF-ChIP signals (Fig. 5F). Together, these results indicate that it may not be required to reposition or deplete nucleosomes for factor binding.

Additional computer simulations confirmed our *in vivo* observations (Fig. 5G). We introduced a nucleosome with an enhanced probability of an intra-nucleosome cleavage (*p*_*cut*_(*nuc, fragile*) = 4 · *p*_*cut*_(*nuc*)), surrounded by two hyper-accessible inter-nucleosome cleavage sites (*p_cut_(linker*, *hyper-accessible) =* 1.2 · *p*_*cut*_(*linker*)) to our *in silico* model. This small adaptation was sufficient to reproduce MNase-sensitive nucleosomes and local hyper-accessible sites observed *in vivo*.

## Discussion

In the present study, we used differential MNase treatment of cellular chromatin and sequencing of the released mono- and di-nucleosomal DNA to study the architecture of the packaged genome. Our results reveal that after normalizing for the genomic GC content, no domains with increased or decreased DNA accessibility could be detected. In the size range of a few kilo base pairs to mega base pairs the DNA is similarly accessible to MNase hydrolysis, suggesting no differences in the level of chromatin packaging. Characterizing and correlating epigenomic features such as histone marks and functional DNA elements with MNase accessibility also did not uncover significant overlaps in altered MNase sensitivity with the respective chromatin marks. Taken together, our data suggests that chromatin is not organized into hierarchical layers of DNA packaging that would progressively decrease the accessibility of DNA.

Our results confirm microscopy studies, revealing the absence of regular nuclear patterns that could be interpreted as higher-order structures of chromatin. McDowell and colleagues visualized a homogenous grainy texture of chromosomes, interpreted as 11 nm fibres, lacking areas of further DNA compaction (McDowall et al. 1986). Additionally, a recent study showed evidence for the lack of domains with distinct higher-order structures of chromatin (Ou et al. 2017). This ChromEMT method beautifully showed that irregular chromatin polymers, with a diameter of 5 to 24 nm, exist throughout the cell cycle. The irregular polymers are present at higher or lower concentrations within different nuclear areas and are highly concentrated within the mitotic chromosomes. In addition to changes in polymer concentration, chromatin did not form specific domains with regular folding in the nucleus (Ou et al. 2017), suggesting the absence of defined higher structures of chromatin. Our study further shows that also within domains of specific epigenetic modifications or genomic functions, no relative difference in DNA accessibility can be detected. This gives support for a model in which, for example euchromatin and heterochromatin domains, with distinct transcriptional activity, do not differ in DNA accessibility. Given the study by Ou and colleagues, which suggests that chromatin domains differ in their local concentration of chromatin (Ou et al. 2017), and the fact that MNase is a small endonuclease, we assume that the enzyme diffuses equally well into all domains of high and low chromatin concentration within the cell. A lack of qualitative differences in MNase dependent cleavage differences implies that there is no distinct structural organization correlating with chromatin density. This finding indicates that the accessibility of DNA to small sequence specific DNA binding factors is similar throughout the genome, arguing against chromatin folding as a mechanism to regulate DNA accessibility and gene activity (Fig. 5H).

A lack of defined chromatin structure raises the question of how chromatin can fulfil its regulatory function and establish defined compartments within the cell. Our data is in good agreement with the polymer melt model presented by Maeshima and colleagues using fluorescently tagged histones and cryo-electron microscopy to study the chromatin architecture (Eltsov et al. 2008; Hihara et al. 2012). The concept of a polymer melt-like structure suggests that the nucleosome fibres are constantly moving and rearranging. high local nucleosome concentrations and interdigitating protein complexes would compete with the formation of regular higher-order structures (Hansen et al. 2017). Our MNase approach ruled out the possibility that the large tag of the fluorescently tagged histone would interfere with the regular folding of chromatin. Taken together, these studies suggest the absence of regular 30 nm fibres and additional higher order structures. However, the ChromEMT study showed, by maintaining the 3D structure of the nucleus, that different parts of the nucleosomal arrays do not directly interact or interdigitate, thereby disturbing the regular path of the fibres (Ou et al. 2017). The fibres are clearly separated in space and theoretically, the isolated fibres should be able to form intra-nucleosomal interactions and regular higher order structures. The absence of fibre contacts suggests that other mechanisms hinder the proper folding of chromatin, which is observed in *in vitro* experiments (Pepenella et al. 2014; Zhu and Li 2016; Schalch et al. 2005; Robinson et al. 2006). We recently showed that chromatin-associated RNA plays an important role in maintaining chromatin accessible. RNase treatment of intact nuclei resulted in the compaction of chromatin and a dramatic reduction in MNase accessibility (Schubert et al. 2012). We concluded that chromatin bound RNA would inhibit the folding of chromatin into defined higher-order structures. With our new study, we would extend this interpretation: We suggest that chromatin-associated RNAs may also regulate the local chromatin density in the cell and more generally interfere with the regular folding of the chromatin fibre. Negatively charged RNA molecules could regulate local chromatin density by a simple charge repulsion mechanism and maintain the nucleosomal arrays at distance and dynamic. Regions of distinct chromatin density, as revealed by ChromEMT, may represent different RNA pools or RNA concentrations associated with chromatin.

Bringing together the irregular, non-structured nature of cellular chromatin, the binding of RNA, the local differences of chromatin density and the characterized epigenomic domains (Roadmap Epigenomics Consortium et al. 2015), a previously described buoy model offers an attractive explanation for the regulation of DNA-dependent processes in the cell (Maeshima et al. 2015). It proposes that the regulation of chromatin domains does not occur at the level of regular structure, but on the level of chromatin density. Small regulatory DNA binding factors, in the size range of the enzyme MNase, would be capable to access dense chromatin domains without hindrance. However, large multi-protein complexes such as the replication and transcription machinery would be expelled from these domains. For example, pioneering transcription factors could migrate into chromatin domains, identify regulatory regions and bind to them. These sites would now, due to transcription factor recruitment and the associated changes in the physicochemical properties of this region, similar to a buoy show up on the surface of the chromatin domain. At the surface of the dense chromatin domain, this genomic region could now be integrated into the large replication or transcription factories. Molecular sieves could have regulatory potential in combination with local, accessible regions, however, further studies are needed to test these models.

A recent chemical map of modified histone H4 crosslinking to DNA in mouse ES cells revealed a broader binding of histone H4 than expected (Voong et al. 2017). Nucleosome free regions (NFR) at active promoters, enhancers and transcription termination regions were defined by their accessibility to MNase and suggested to have nucleosomes replaced by DNA binding factors (Hughes and Rando 2014). However, Voong and colleagues were able to detect histone H4 binding to these sites, implying the presence of sensitive or altered nucleosomes being bound to these sites (Voong et al. 2017). These authors even claim that histone occupancy is higher at active genes, as more H4 can be crosslinked. Our study, using low MNase concentrations, is now clearly revealing nucleosome-sized DNA fragments at the previously denominated NFRs. Taken together, we suggest that MNase sensitive nucleosomes locally occupy these active genomic regions and probably co-occupy these sites with DNA binding factors such as CTCF. Nucleosomal MNase sensitivity could be a result of a) active nucleosome remodelling and high turn-over, as catalysed by the remodelling enzyme RSC (Kubik et al. 2015); b) the active incorporation of histone variants such as H3.3 and H2A.Z that could destabilize promoter nucleosomes (Zlatanova and Thakar 2008; Bruce et al. 2005); c) histone modifications that increase DNA breathing and octamer stability (Ausio and van Holde 1986; Poirier et al. 2008); or d) being an effect of an altered nucleosome structure, as described by Allfrey and colleagues (Prior et al. 1983). The nature of these unstable nucleosomes still must be determined biochemically, but intriguingly they appear as individual particles localized in between non-sensitive histone octamers.

## Methods

### Differential MNase hydrolysis and DNA isolation

Human adenocarcinoma cells of the HeLa line were cultured in D-MEM medium supplemented with 10% FCS. Cells were incubated at 37°C with 5% CO_2_ on 15cm cell culture dishes. Cells were grown to 70% to80% confluence and dishes were washed once with 5 ml PBS. Cells were incubated for 3 min with 3 ml of permeabilisation buffer (15 mM Tris/HCl pH 7.6, 300 mM saccharose, 60 mM KCl, 15 mM NaCl, 4 mM CaCl_2_, 0.5 mM EGTA, 0.2% (v/v) NP40 and fresh 0.5mM 2-mercaptoethanol) including 100 to 2000 units of MNase (Sigma) (Stewart et al. 1991). The DNA hydrolysis reaction was stopped by the addition of 3 ml of stop buffer (50mM Tris/HCl pH8, 20mM EDTA and 1% SDS). RNA was digested by the addition of 250 μg RNase A and incubation for 2h. Subsequently 250 μg of proteinase K were added to the culture dishes and incubated over night at 37°C. Fragmented DNA was purified by ammonium acetate and ethanol precipitation. The MNase digestion pattern was analyzed on 1.3% agarose gels and visualized after ethidium bromide staining.

MNase digestion reactions were loaded on preparative agarose gels (1.1% agarose) electrophoresed and the mono-, di- and tri-nucleosomal DNA were excised after ethidium bromide staining. DNA was purified by electroelution using the Electro-Eluter 422 (BioRad). DNA was precipitated and re-analyzed by agarose gel electrophoresis.

### 2D and 3D fluorescence in situ hybridisation

The isolated tri-nucleosomal DNA isolated from low- and high-MNase samples were labelled with the Prime-it Fluor Fluorescence Labelling Kit (Stratagene) according to manufacturers’ protocol. 1 μg of total DNA was labelled with fluor-12-dUTP or Cy3-dUTP using the random priming protocol. 2D FISH experiments were performed on metaphase spreads of normal human lymphocytes, as described (Cremer et al. 2008). Cot1 DNA (10 ug) was used in the hybridisation reactions to suppress the signals of repetitive DNA regions.

3D FISH experiments were performed as described (Cremer et al. 2008). IMR90 human diploid fibroblast cells were fixed and 3D FISH experiments were performed. Confocal microscopy and image analysis was done after 3D FISH experiments as follows: A series of optical sections through 3D-preserved nuclei were collected using a Leica TCS SP5 confocal system equipped with a Plan Apo 63-/1.4 NA oil immersion objective and a diode laser (excitation wave length 405 nm) for DAPI, a DPSS laser (561 nm) for Cy3 and a HeNe laser (633 nm) for Cy5. For each optical section, signals in different channels were collected sequentially. Stacks of 8-bit grey-scale images were obtained with z-step of 200 nm and pixel sizes 30 - 100 nm depending on experiment.

### DNA sequencing, alignment and data processing

Sequencing libraries of the high and low MNase treated nucleosomal DNA (mono- and di-nucleosomal DNA) were prepared using the NEBNext DNA Library Prep Master Kit (New England Biolabs, Ipswich, USA) and paired-end (2 × 50 bp) sequenced on a HiSeq2000 platform (Genecore, EMBL, Heidelberg). The sequenced paired-end reads were mapped to the UCSC human genome version 37 (hg19) using the local alignment option of `bowtie2` with following parameters -- `very-sensitve-local` and --`no-discordant` (Langmead and Salzberg 2012). Aligned reads were filtered for mapping quality (`MAPQ` > 20) and further processed using `SAMtools` (Li et al. 2009) and `BEDtools` (Quinlan and Hall 2010). Fragments with a size of 140 - 200 bp were considered to originate from mono-nucleosomal protected DNA and fragments with a size of 250 - 500 bp as di-nucleosomal respectively. All downstream analyses were performed on the filtered and size-selected fragments.

### Genome wide nucleosome maps

Genome wide nucleosome occupancy maps were generated using the `danpos`. `py` script from the `DANPOS2` toolkit (Chen et al. 2013) in `dpos` mode. Here, reads were extended to 70 bp following the centering to the fragment midpoint, to obtain focused mono-nucleosomes (150 bp for di-nucleosomes). Clonal reads were filtered using the `parameter u 1E-2` and the total number of fragments was normalized to 200 million fragments for each sample.

Obtained nucleosome occupancy maps were further processed using `Control-FREEC` v8.7 to call copy number alterations (CNV) in HeLa cells (Boeva et al. 2012). `Control-FREEC` was ran on single-end high-throughput sequencing data of sonicated chromatin in HeLa cells, which were annotated as above, except that reads were not filtered without matching read-pair as unbiased control with following settings: `ploidy = 3, window = 500,000, step = 50,000, minMappabilityPerWindow = 0.80, breakPointThreshold = 3`. The algorithm also accounts for the underlying GC content and mappability of the genome. Therefore, the provided mappability track covering up to 2 mismatches for 50 bp single-end reads (available at http://boevalab.com/FREEC/) and a GC profile consisting of 500 kb sliding windows with 50 kb steps have been integrated in the analysis. Finally, nucleosome occupancy maps have been normalized by the reported CNVs.

### Differential MNase analysis

The complete human genome (hg19) was segmented in non-overlapping windows of a constant size (ranging from 250 bp – 1 Mb). The average nucleosome profile level of each MNase condition, the average GC content and mappability score which was based on CRG alignability track for 100mers (Derrien et al. 2012), was calculated for each window. Windows lacking mapped reads or exhibiting an average mappability score below 0.9, were excluded from further analyses. Linear regression analysis was carried out in the `R` environment (R Core Team 2014) applying the implemented linear regression function (`lm`). Extreme values exhibiting a high influence on the regression analysis were filtered using a Cook’s distance threshold of 0.001. Adjusted R^2^ values reported by `lm` were used to assess the dependency of the tested features. Density plots were drawn with the `heatscatter` function implemented in the `R` package `LSD` (Schwalb et al. 2018).

### GC content and di-nucleotide frequency of nucleosomal fragments

The average GC content and dinucleotide frequency around the borders of mapped fragments were calculated using the `makeTagDirectory` script from the `HOMER` software package (Heinz et al. 2010). To calculate the dinucleotide frequency around the midpoints of mapped fragments, properly aligned read-pairs were converted to a bed format comprising start and end positions of the fragments using `BEDtools`. The dinucleotide frequency was obtained by applying the `annotatePeaks`.`pl` script of the HOMER software package with the parameters -`hist 1, − di -size −300,300`.

### Nucleosome prediction, average nucleosome occupancy profiles and heatmaps

The `dpos` script from the `DANPOS2` toolkit was applied to the normalized nucleosome occupancy profiles with default settings in order to predict nucleosome positions (Chen et al. 2013). Average nucleosome occupancy profiles and/or heatmaps around factor binding sites, transcription start sites or DNase I HS were generated using the `DANPOS2` script `profile`.

### GC normalization and detection of local differential chromatin accessibility

A locally weighted scatterplot smoothing (LOESS) was applied to correct for the inherent GC bias in the nucleosome occupancy profiles. Therefore, the average nucleosome profile level and GC content within 250 bp non-overlapping windows were calculated and filtered as described above. Additionally, windows exhibiting an extreme GC content (< 30% or > 70%) were filtered. The `loess` function implemented in the R environment was applied to estimate the influence of GC content on high or low MNase digestions. The nucleosome occupancy profile in the 250 bp windows was corrected by the LOESS analysis determined correction factor. Local differences of GC-normalized low and high MNase nucleosome occupancy profiles were determined using the dpeak algorithm of the `DANPOS2` toolkit with the parameters --`peak_width 70, -p 1E-2`. Differentially enriched peaks were selected by a false discovery rate (FDR) below 0.05 and were termed as MNase hyper-accessible (enriched in low MNase) or MNase in-accessible (enriched in high MNase).

### Simulation of MNase digestions

Simulations were conducted on an artificially generated nucleosome fibre comprising of 50 nucleosomes of 150 bp in size, regularly spaced onto a 9000 bp long DNA strand. One potential MNase cleavage site was assigned to each linker DNA and each nucleosome. First, an index containing the positions of the potential cleavage sites *PosCut* was created. Cleavage sites *posCuti* were assigned to every 90 bp ((150 bp nucleosomal DNA + 30 bp linker DNA)/2) of the 9000 bp long strand, representing alternating linker *posCut(linker)*_*i*_ and nucleosome *posCut(nuc)*_*i*_ cleavage sites. Every second *posCut(nuc)* was shifted 5 bp downstream, consequently *posCut(nuc)_*i+1*_* − *posCut(nuc)_i_* = 180 ± 5 bp and *posCut(linker)*_*i+1*_ − *posCut(linker)*_*i*_ = 180 bp.

In total, this generated an index of the cleavage positions along the simulated DNA strand: *PosCut=(posCut(linker)_1_ = 0, posCut(nuc)_1_ = 85, posCut(linker)_2_ = 180, posCut(nuc)_2_ = 270,…, posCut(nuc)_50_ = 8910, posCut(linker)_51_ = 9000)*.

In the next step, the *posCut(nuc)*_*i*_ and *posCut(linker)*_*i*_ values were sampled according to their associated probability of an intra-nucleosomal MNase cleavage event, namely *p*_*cut*_(*nuc*) or inter-nucleosomal MNase cleavage event *p*_*cut*_(*linker*). Initially, we chose the probability of *p*_*cut*_(*linker*) = 10 × *p*_*cut*_(*nuc*) according to *in vitro* studies (Cockell et al. 1983). Subsequent values were adapted as indicated in the text. From this distribution, *n* values *posCut(nuc)_i_* or *posCut(linker)_i_* were randomly drawn without replacement, simulating *n* cutting events on one template. For high MNase simulations, *n* = 70 cutting events were chosen per template and *n* = 20 for low MNase simulations. The corresponding cutting positions *posCol* were sorted in ascending order: *PosCol = (posCol_1_* < *posCol_2_ <…< posCol_n_)*.

Neighbouring positions *posCol_j_* were assembled into fragments *fr(posCol_i_, posCol_i+1_)* and fragments of 180 bp length were stored, since they represent the simulated mono-nucleosomes. This process was iterated until one million fragments of 180 bp length were sampled. Finally, the nucleosomal fragments were trimmed by 30 bp from both sides to centre the nucleosome positions and the coverage over the simulated strand was plotted.

### External data sets used in this study

DNase-seq (ENCSR959ZXU), CTCF binding sites (ENCSR000CUB), H3K27ac-ChIP (ENCFF311EWS), H3K27me3-ChIP (ENCFF958BAN), and polyA mRNA RNA-seq (ENCSR000EYQ) tracks were downloaded from the ENCODE repository (ENCODE Project Consortium et al. 2012; Sloan et al. 2016). Chromatin state annotation for HeLa cells (E115, 15 core marks) was obtained from the roadmap epigenomics project (Roadmap Epigenomics Consortium et al. 2015). Enhancer site analysis was carried out using the predefined HeLa enhancer sets from (Chen et al. 2016b). CRG mappability (align 100 mer) track was downloaded from the UCSC browser (http://hgdownload.soe.ucsc.edu/goldenPath/hg19/encodeDCC/wgEncodeMapability/).

## Data access

MNase-seq data has been submitted to NCBI Gene Expression Omnibus (GEO; http://www.ncbi.nlm.nih.gov/geo/) under accession number GSE100401.

## Acknowledgements

We thank Thomas Cremer and Irina Solovei for their help in imaging.

